# Structures of human cytoplasmic dynein in complex with the lissencephaly 1 protein, LIS1

**DOI:** 10.1101/2022.10.08.511426

**Authors:** Janice M. Reimer, Morgan E. DeSantis, Samara L. Reck-Peterson, Andres E. Leschziner

## Abstract

The lissencephaly 1 protein, LIS1, is mutated in type-1 lissencephaly and is a key regulator of cytoplasmic dynein-1. At a molecular level, current models propose that LIS1 activates dynein by relieving its autoinhibited form. We recently reported a 3.1Å structure of yeast dynein bound to Pac1, the yeast homologue of LIS1, which revealed the details of their interactions (Gillies et al., 2022). Based on this structure, we made mutations that disrupted these interactions and showed that they were required for dynein’s function in vivo in yeast. We also used our yeast dynein-Pac1 structure to design mutations in human dynein to probe the role of LIS1 in promoting the assembly of active dynein complexes. These mutations had relatively mild effects on dynein activation, suggesting that there may be differences in how dynein and Pac1/LIS1 interact between yeast and humans. Here, we report cryo-EM structures of human dynein-LIS1 complexes. Our new structures reveal the differences between the yeast and human systems and provide a blueprint to disrupt the human dynein-LIS1 interaction more accurately. Our new structures also allow us to map type-1 lissencephaly disease mutations, as well as mutations in dynein linked to malformations of cortical development/ intellectual disability, in the context of the human dynein-LIS1 complex.

## Introduction

Cytoplasmic dynein-1 (dynein here) is a conserved microtubule-based molecular motor. In humans, dynein moves dozens of distinct cargos towards the minus ends of microtubules (Reck-Peterson et al., 2018), while in yeast dynein has a single known role in aligning the mitotic spindle (Markus et al., 2020). Active dynein complexes are composed of one or two dimers of dynein, the dynactin complex, and an activating adaptor (Grotjahn et al., 2018; McKenney et al., 2014; Schlager et al., 2014; Urnavicius et al., 2018). In cells, dynein dimers are thought to exist primarily in an autoinhibited form (Amos, 1989; Torisawa et al., 2014; Zhang et al., 2017), which must be relieved for cargo movement. Recent work has shown that LIS1 has a conserved role in relieving dynein autoinhibition (Elshenawy et al., 2020; Gillies et al., 2022; Htet et al., 2020; Qiu et al., 2019). At a functional level, Lis1 is a dimer of two β-propellers (Kim et al., 2004; Tarricone et al., 2004). *LIS1* was originally described as the gene mutated in patients with type-1 lissencephaly (Parrini et al., 2016; Reiner et al., 1993). Later work linked *LIS1* to the dynein pathway (Sasaki et al., 2000; Smith et al., 2000; Tai et al., 2002; Xiang et al., 1995). Mutations in the dynein motor-containing heavy chain (*DYNC1H1*) have also been linked to malformations of cortical development (Lipka et al., 2013; Parrini et al., 2016). Despite the importance of LIS1 in understanding these human diseases, no three-dimensional structures of a human dynein-LIS1 complex have been reported.

Cytoplasmic dynein is a member of the AAA+ (ATPase associated with various cellular activities) family of proteins. Unlike most members of the AAA+ family, which are oligomers, the AAA+ domains in dynein’s “heavy chain” are fused into a single polypeptide with each of its six AAA+ domains having diverged over time (Canty and Yildiz, 2020). Four of dynein’s AAA+ domains can bind ATP (AAA1-AAA4), and three hydrolyze it (AAA1, AAA3, and AAA4). AAA5 and AAA6 have diverged enough to no longer be able to bind nucleotides (Schmidt and Carter, 2016). Dynein’s heavy chain can be divided into several elements with specific functions (Figure 1A). At its amino-terminus, the “tail” is responsible for dimerization, and is the site for binding of several accessory subunits. The tail is followed by the “linker”, a mechanical element that undergoes conformational changes, bending at a “hinge” in response to the nucleotide state of dynein’s AAA+ “ring” to drive movement. Two elements protrude from dynein’s ring: the “stalk”, a long antiparallel coiled-coil that connects the ring to the microtubule-binding domain (MTBD), and the “buttress”, a short antiparallel coiled-coil that couples conformational changes in the ring with conformational changes in the MTBD by altering the register between the two alpha helices in the stalk (reviewed in Cianfrocco et al., 2015). Dynein’s AAA+ ring mainly exists in one of two conformations driven by the nucleotide state of its AAA+ domains: an “open” conformation coupled to high affinity for the microtubule, and a “closed” conformation that leads to low affinity for the microtubule (Schmidt et al., 2015, 2012). In dynein’s autoinhibited state, called “Phi” due to its resemblance to the Greek letter, the two heavy chains come together face to face, thus pointing in opposite directions and preventing the motor from engaging microtubules in a manner that would allow it to walk (Zhang et al., 2017).

**Figure 1.**
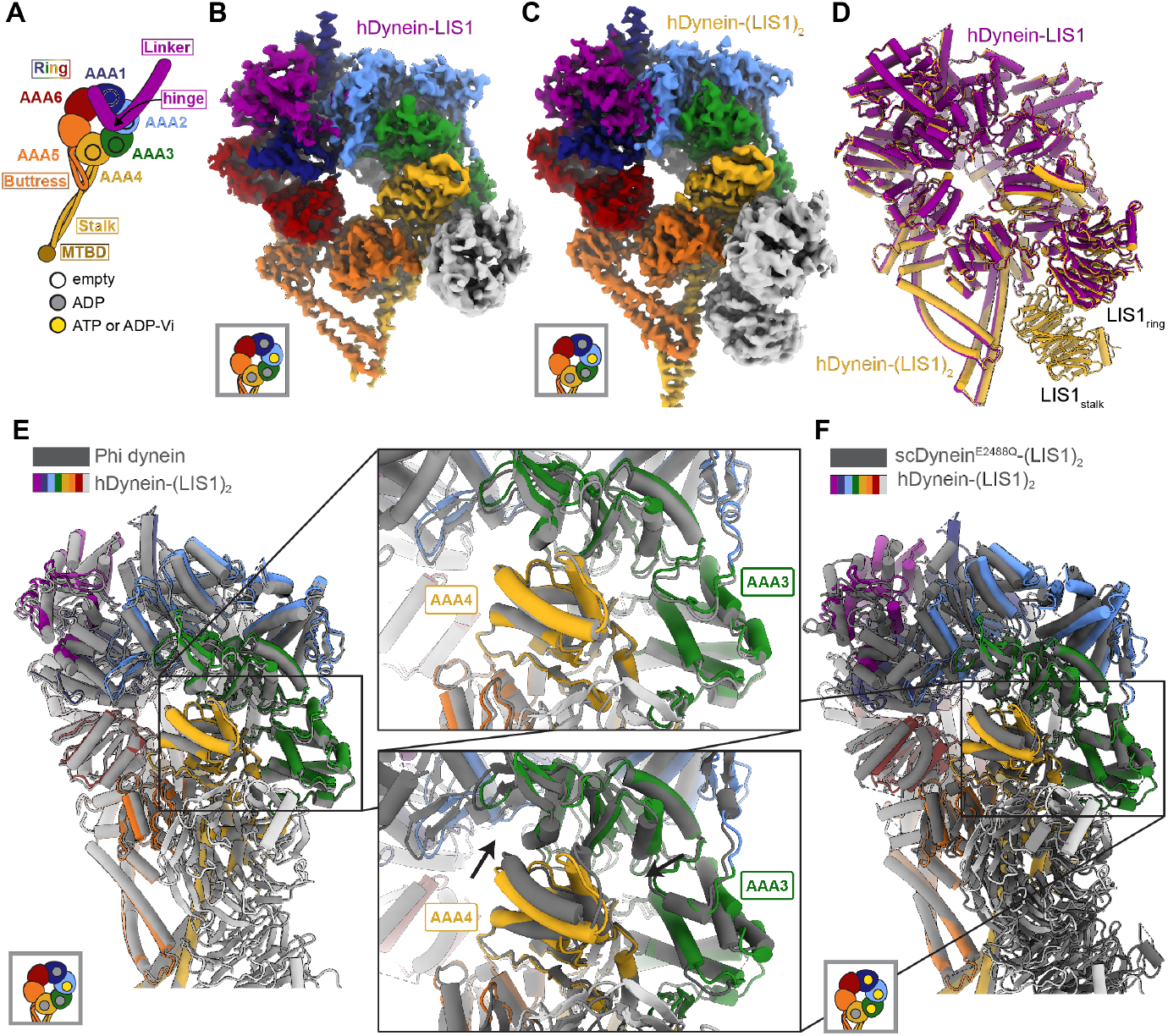
Structures of human dynein bound to LIS1. **(A)** Cartoon schematic of dynein showing domain organization. The names of the major structural elements in dynein are indicated inside boxes. MTBD: MicroTubule Binding Domain. The four AAA+ domains that can bind nucleotide are indicated with the black circles. The color coding used throughout the figures to indicate their nucleotide state is shown below dynein’s cartoon. **(B, C)** Cryo-EM maps of human dynein bound to (B) one (dynein-LIS1) or (C) two (dynein-(LIS1)_2_) LIS1 β-propellers. **(D)** An overlay of the two human dynein-LIS1 structures solved here. **(E)** An overlay of the human Phi dynein (PDB 5NVU) and the human dynein-(LIS1)_2_ structures, aligned on AAA4. The inset shows that the conformation of AAA3 and AAA4 are the same. **(F)** An overlay of the yeast dynein-(Pac1)_2_ (carrying a point mutation at E2488Q; PDB 7MGM) and the human dynein- (LIS1)_2_ structures, aligned on AAA4. The inset shows there is a slight difference in AAA4 relative to AAA3 between the two structures.

We recently reported a 3.1Å structure of the yeast dynein-Pac1complex (Gillies et al., 2022). In this structure, dynein is present as a single motor domain bound to two Pac1 β-propellers. This was the first high-resolution structure of a complex between dynein and Pac1 and revealed a number of interactions that earlier, lower resolution maps of the complex had failed to identify (DeSantis et al., 2017; Huang et al., 2012; Toropova et al., 2014). By designing mutants based on this structure, we showed that binding of Pac1 to dynein, either to its AAA+ ring (site_ring_) or its stalk (site_stalk_), and the interaction between the two Pac1 β-propellers were essential for dynein’s function in yeast in vivo (Gillies et al., 2022). We also used our model of the yeast dynein-Pac1 complex to identify mutations in human dynein to probe the role of LIS1 in relieving human dynein autoinhibition (Gillies et al., 2022). These mutations had relatively mild effects on dynein activation, suggesting that there may be differences in how dynein and LIS1 interact between yeast and humans that our modeling did not capture.

Here, we set out to determine a high-resolution structure of the human dynein-LIS1 complex. Previously we had obtained 2D class averages from cryo-EM datasets of human dynein in the presence of LIS1 showing that LIS1 binds dynein simultaneously at site_stalk_ and site_ring_, as was the case in yeast (Htet et al., 2020). The human and yeast structures appeared very similar at the level of the 2D class averages (Htet et al., 2020). We now report 3D cryo-EM structures of human dynein bound to one and two human LIS1 β-propeller domains. We show that there are differences in how human LIS1 binds dynein relative to its yeast counterpart. We compare the human and yeast structures, focusing on the interactions between dynein and LIS1/Pac1 at site_stalk_ and site_ring_ and the LIS1-LIS1/Pac1-Pac1 interaction. Overall, our work provides a model for how human LIS1 interacts with human dynein. Importantly, our structures also allow us to map the type-1 lissencephaly disease mutations, as well as dynein disease mutations, including those that cause malformations of cortical development/intellectual disability, in the context of dynein’s interaction with LIS1.

## Results and Discussion

### Structures of human dynein bound to LIS1

We revisited our cryo-EM datasets of human dynein and LIS1 (Htet et al., 2020) and, with additional processing, solved the structures of human dynein bound to one and two LIS1 β-propellers to 4.0 Å and 4.1 Å, respectively (Figures 1B and 1C; Figure 1 – figure supplement 1; Table 1). In both structures, dynein is in the closed ring conformation and the linker domain is disordered before the hinge region. The conformation of dynein is the same regardless of whether LIS1 is bound only at site_ring_ or both at site_ring_ and site_stalk_ (Figure 1D). The closed state of the motor domain seen in our structures is the same as that observed in the autoinhibited Phi conformation of human dynein (Figure 1E) (Zhang et al., 2017).

We prepared our samples with ATP and vanadate included in the buffer. Hydrolysis of ATP by dynein in the presence of vanadate leads to the formation of ADP-V_i_, a post-hydrolysis ADP.P_i_ analogue. Based on the map density, ADP is bound to AAA1 and AAA4, while AAA2 contains either ATP or ADPV_i_ (Figure 1 – figure supplement 2). The nucleotide state of AAA3 is unclear from the density, but we chose to model ADP (Figure 1 – figure supplement 2) as AAA3 has the same conformation as the ADP-bound AAA3 domain observed in the structure of human Phi dynein (Figure 1E), while the conformation of AAA3 in yeast dynein (carrying a point mutation, E2488Q, at AAA3) is different when bound to ATP (Figure 1F).

### LIS1 binding to dynein at site_ring_

Human LIS1 binds dynein in a manner similar to that of yeast dynein at site_ring_. At site_ring_, the main contact between LIS1 and dynein involves the same AAA4 helix used by yeast dynein, as well as the same AAA5 loop and the loop bridging AAA3-AAA4 (Figure 2A; Video 1). We previously showed that the AAA4 and AAA5 interactions with yeast Pac1 are important for dynein regulation (Gillies et al., 2022). There is a minor rotation in human LIS1_ring_ relative to yeast Pac1_ring_ that causes a slight shift in how LIS1 interacts with dynein (Figures 2B and 2C; Video 1). Despite these changes, the interfaces between dynein and LIS1 we saw in our yeast dynein-Pac1 structure are maintained; the AAA5 loop appears to make a small compensating shift to preserve its contact with LIS1 (Figure 2B). Additionally, the placement of LIS1 at site_ring_ is the same whether one or two LIS1s are bound to human dynein (Figure 1D).

**Figure 2.**
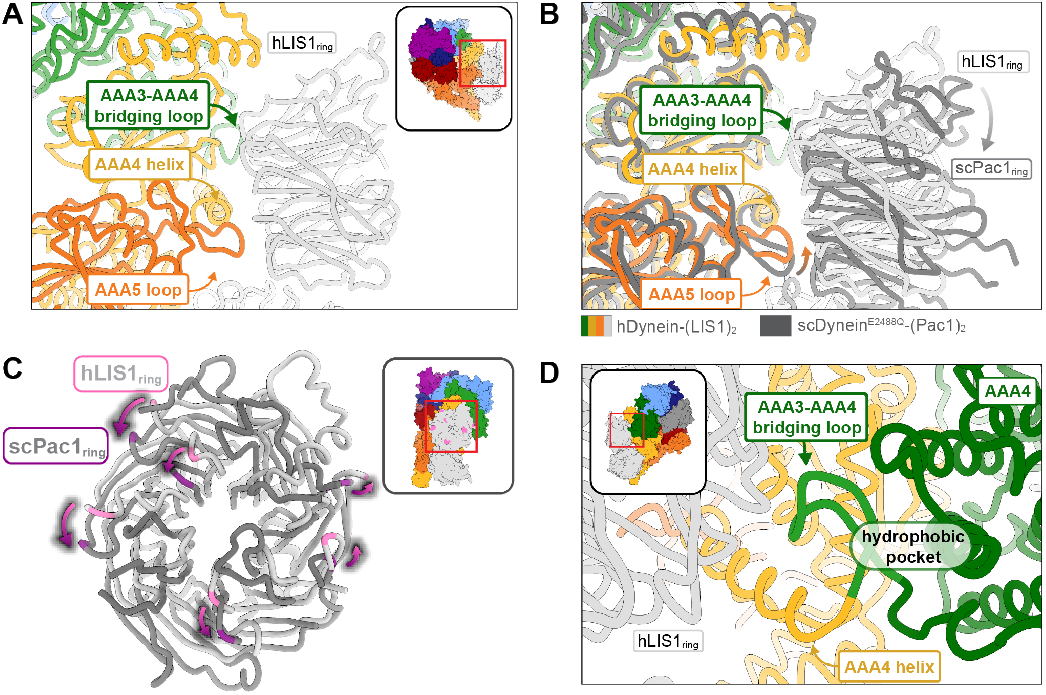
Structure of LIS1 binding to dynein at site_ring_. **(A)** LIS1 at site_ring_ interacts with dynein via the AAA3-AAA4 bridging loop, a AAA4 helix and a AAA5 loop. **(B)** An overlay of the human and yeast dynein structures bound to LIS1/Pac1, aligned on AAA4 (human, light grey; yeast, dark grey). **(C)** LIS1 (light grey) and Pac1 (dark grey) from panel are viewed facing the β propeller, with dynein removed for clarity. This panel shows the rotation, highlighted by the purple markers and arrows of LIS1 relative to Pac1 at site_ring_. **(D)** The AAA3-AAA4 bridging loop contacts LIS1 and preserves a hydrophobic pocket in AAA4.

The contact between the AAA3-AAA4 bridging loop and LIS1 (Figure 2D) is the pivot point about which the position of LIS1_ring_ rotates between the human and yeast systems (Video 1). Given the conservation of this interaction between the two systems, an intriguing possibility is that this contact relays information about the nucleotide state of AAA3, a regulatory site in dynein (DeSantis et al., 2017; Dewitt et al., 2015; Dutta and Jana, 2019; Nicholas et al., 2015; Qiu et al., 2021), to LIS1/Pac1. The loop forms a hydrophobic pocket with the small domain of AAA3 (AAA3_S_), and nucleotide-induced conformational changes in AAA3 may cause this loop and AAA3_S_ to shift together (Figure 2D). The bridging loop could thus act as a tether between LIS1 and AAA3 to impede the release of ADP from AAA3.

### LIS1 binding to dynein at site_stalk_

At site_stalk_, LIS1 interacts with dynein at both the CC1 helix in the stalk (the helix leading from dynein’s ring to the MTBD) and at a loop in AAA4 (residues 3112-3119) (Figure 3A). Human LIS1 is pivoted around the stalk helix relative to yeast Pac1 to a larger extent than at site_ring_ (Figure 3B and Video 1). We originally used a yeast dynein^E3012A Q3014A N3018A^ mutant (“dynein^EQN^”) (DeSantis et al., 2017) to probe the importance of the Pac1-stalk interaction (Figure 3C). These mutation sites were chosen based on sequence conservation and low resolution cryo-EM models. Comparing our yeast dynein-Pac1 structure to our new human dynein-LIS1 structure shows that the EQN triad residues are shifted relative to where we had modeled them (Video 1), providing an explanation for the modest phenotype we observed when we mutated these residues in human dynein (Gillies et al., 2022).

**Figure 3.**
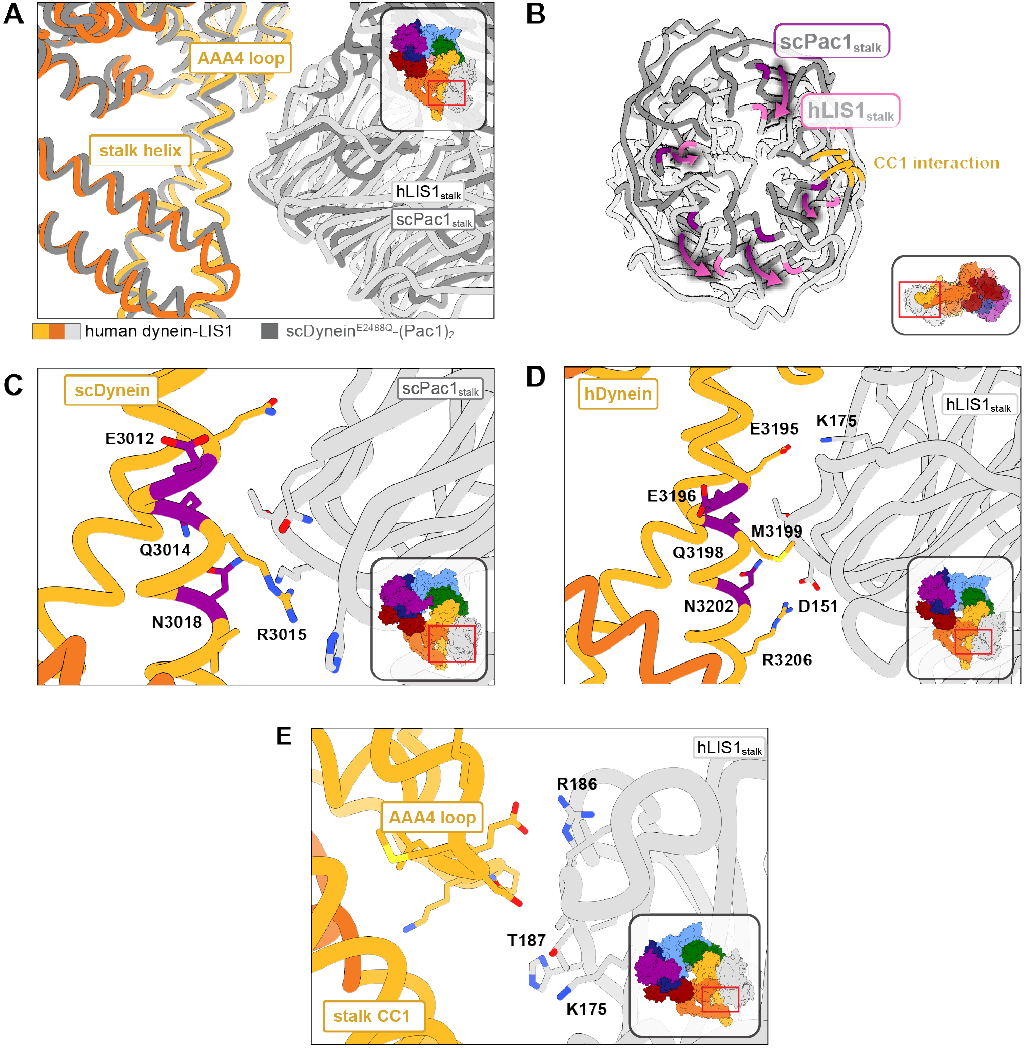
Structure of LIS1 binding to dynein at site_stalk_. **(A)** An overlay of human and yeast dynein bound to LIS1/Pac1, aligned on AAA4. **(B)** LIS1 (light grey) and Pac1 (dark grey) from panel (B) are viewed facing the β-propeller, with dynein removed for clarity. This panel shows the rotation, highlighted by the purple markers and arrows of LIS1 relative to Pac1 at site_stalk_. The area where LIS1/Pac1 interacts with dynein’s CC1 stalk helix is shown in yellow. **(C)** The yeast dynein-Pac1_stalk_ interaction. **(D)** The human dynein-LIS1_stalk_ interaction. **(E)** The AAA4 loop – LIS1_stalk_ interaction.

The structure of yeast dynein-Pac1 (Gillies et al., 2022) showed that N3018 is the only residue in the EQN triad that forms a hydrogen bond with Pac1, although Q3014 may act to stabilize N3018 by forming a small hydrogen bonding network. R3015 and Q3011 form two additional hydrogen bonds with the backbone of Pac1. Hence, a yeast dynein^R3015 Q3011 N3018^ mutant may be better than the original dynein^E2012 Q3014 N3018^ mutant used to disrupt Pac1 regulation at site_stalk_. Similarly, the equivalent EQN triad in human dynein, E3196, Q3198, and N3202, follows the same interaction pattern, where Q3198 interacts with N3202, but only N3202 hydrogen bonds with the backbone of LIS1 (Figure 3D). R3206 is in position to form a salt bridge with LIS1 D151, which is made possible by the rotation of LIS1 in the human structure relative to the yeast one. We predict that point mutations at M3199A, N3202A, and R3206A in human dynein would disrupt LIS1 regulation at site_stalk_ to a greater extent than dynein^E3196 Q3198 N3202^. A septuple mutant we designed in human dynein, dynein^K2898A E2902G E2903S E2904G E3196A Q3198A N3202A^, which comprised the EQN mutations, was still capable of binding LIS1 (Gillies et al., 2022).

Previously we showed that yeast Pac1^S248Q^ acts as a separation-of-function mutant that disrupts the regulation of Pac1 at site_stalk_ (Gillies et al., 2022). In the human structure, the LIS1_stalk_-AAA4 loop interaction is primarily mediated through backbone interactions. Based on sequence alignments, T187 in human LIS1 is homologous to yeast S248; however, in our structure T187 faces away from dynein and is poised to hydrogen bond with K175. To make a separation-of-function mutant in human LIS1, the neighboring residue, R186, which extends towards dynein, may serve as a better mutation candidate in future studies (Figure 3E).

### LIS1-LIS1 interaction

The biggest difference between the human dynein-LIS1 and yeast dynein-Pac1 complexes is in the LIS1-LIS1/ Pac1-Pac1 interaction (Video 1). The rotation of LIS1 at site_ring_ and site_stalk_ in the human complex causes the LIS1-LIS1 interface to become significantly smaller, with approximately half the amount of buried surface area (∼301Å^2^) compared to the yeast Pac1-Pac1 interface (∼590Å^2^) (Figure 4A and 4B). However, the chemical nature of the interface is also different: while the yeast Pac1-Pac1 interaction is moderately hydrophobic, the human LIS1-LIS1interface is more electrostatic (Figure 4C and 4D), which may compensate for the smaller surface area.

**Figure 4.**
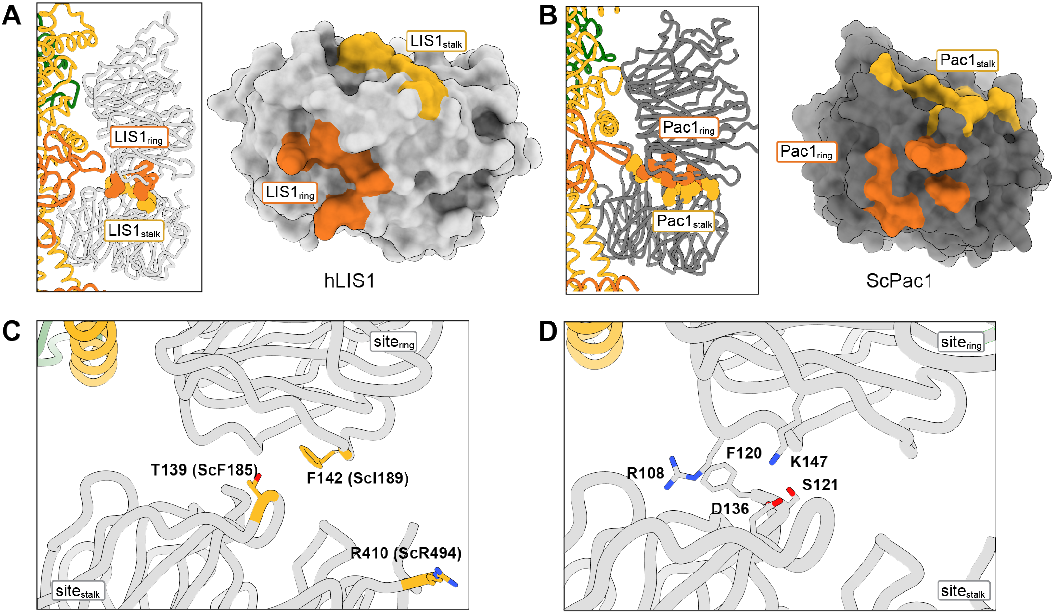
Structure of the LIS1-LIS1 interface. **(A, B)** Residues involved in the LIS1-LIS1 interaction are shown in the context of the dynein-(LIS1/Pac)_2_ structures and mapped onto a surface representation of LIS1 (A) or Pac1 (B). Residues involved in the interaction with site_ring_ (LIS1_ring_) are shown in orange and those involved in the interaction with site_stalk_ (LIS1_stalk_) are shown in yellow. **(C)** The human LIS1-LIS1 interaction does not use residues equivalent to those in the yeast Pac1-Pac1 interaction. **(D)** The human LIS1-LIS1 interface.

The yeast Pac1-Pac1 interface mutations (F189D, I189D, R494A) we previously tested (Gillies et al., 2022) were designed to disrupt the Pac1-Pac1 interface and are not conserved. Based on structure and sequence alignments, the equivalent residues in human LIS1 (T139, F142, R410) do not participate in the LIS1-LIS1 interface and mutating them would likely not have a disruptive effect (Figure 4C). Instead, S121, D136 and K147 may be better candidates to disrupt human LIS1-LIS1 interface (Figure 4D).

### Lissencephaly disease-causing mutations

Lissencephaly is a neurodevelopmental disease caused by mutations in *LIS1* that result in impaired neuronal migration (Parrini et al., 2016; Reiner et al., 1993). Lissencephaly is a disease of haploinsufficiency and the majority of disease-causing mutations in *LIS1* include large deletions or nonsense mutations that lead to truncated products (Cardoso et al., 2002; Haverfield et al., 2009; Lipka et al., 2013; Pilz et al., 1998; Sapir et al., 1999). Mutations are located in both the amino-terminal dimerization domain (LisH) and the WD40 domain (Figure 5A). Missense mutations are less common. Several missense mutations found in the interior of the WD40 domain are part of the DHSW motifs involved in stabilizing the β-propeller fold and are likely to disrupt the structure of the domain. In Video 2 we show the location of those missense lissencephaly mutations where a destabilizing effect was not obvious from an inspection of the structure. In addition to the lissencephaly mutations, Video 2 also shows the location of mutations in LIS1 associated with Miller-Dieker lissencephaly syndrome, subcortical band heterotopia and double cortex syndrome (Haverfield et al., 2009; Pilz et al., 1998; Reiner et al., 1993; Sapir et al., 1999).

**Figure 5.**
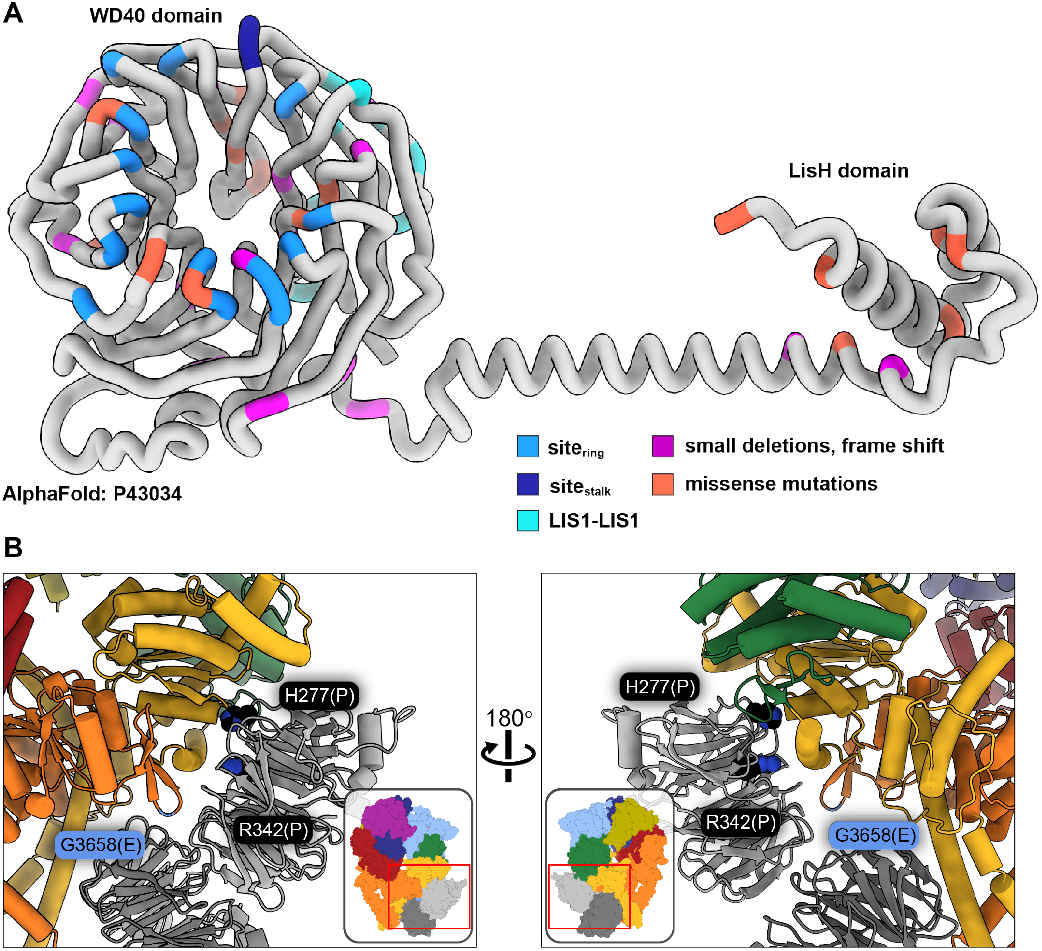
Lissencephaly disease causing mutations. **(A)** AlphaFold (Jumper et al., 2021; Senior et al., 2020) model of full length human LIS1 with residues colored by either interface or lissencephaly mutation. Site_ring_, medium blue; site_stalk_, dark blue; LIS1-LIS1, turquoise; lissencephaly small deletions leading to a frame shift, purple; missense mutations, salmon. **(B)** Two views are shown of disease-linked mutations in dynein located near sites of LIS1 binding. H277P, a lissencephaly mutation, and R342P, a double cortex syndrome mutation, are close to site_ring_. G3658E, associated with intellectual disability, is located at the tip of the AAA5 beta hairpin loop that is part of site_ring_.

Only two known disease-related missense mutations in LIS1 are near a dynein binding site: both H277P, a lissencephaly mutation, and R342P, a double cortex syndrome mutation, are close to site_ring_ (Figure 5B). Although neither amino acid makes a direct contact with dynein, both are involved in hydrogen bonds with nearby residues in the area that comprises the main interface involved in binding to the AAA4 helix at site_ring_. It is likely that local disruption of the structure in the Proline mutants would affect the binding of LIS1 to dynein. H389Y, a subcortical band heterotopia mutation, is located near the LIS1-LIS1 interface. While not part of that interface, H389 makes hydrogen bonds with residues on the same LIS1 β-propeller, including Y137, located in the loop that contains the interface-forming T139.

Video 2 also shows the location of missense mutations in dynein associated with several neurodevelopmental and neurodegenerative disorders: Charcot-Marie-Tooth, spinal muscular atrophy, autism spectrum disorders, and malformations of cortical development/ intellectual disability (Lipka et al., 2013; Reiner et al., 2016; Sabblah et al., 2018; Weedon et al., 2011). One mutation is intriguing in terms of LIS1 regulation of dynein: G3658E, which is associated with intellectual disability (Hertecant et al., 2016). Although G3658 does not interact with LIS1, it is located at the tip of the AAA5 beta hairpin loop that is part of site_ring_ and is likely involved in the formation of the beta hairpin (Figure 5B).

### Conclusions

The cryo-EM structures of human dynein-LIS1 complexes we reported here revealed that while the overall structure of dynein’s interaction with LIS1/Pac1 is conserved from yeast to humans, there are important differences in the specifics of the dynein-LIS1/Pac1 and LIS1-Lis1/Pac1-Pac1 interactions. The data and discussion presented here provide a blueprint to better disrupt the human dynein-LIS1 interface and to map human disease mutations discovered in the future in the context of the human dynein-LIS1 complex.

## Acknowledgements

We thank Eva Karasmanis and Agnieszka Kendrick for editing the manuscript. We thank our funding sources: JMR was a Merck Fellow of the Damon Runyon Cancer Research Foundation, DRG-2370-19; MED was a Jane Coffin Childs Postdoctoral Fellow; SRP’s lab is supported by the Howard Hughes Medical Institute and NIH R35 GM141825; AL’s lab was supported by NIH R01 GM107214 and is now supported by R35 GM145296. We thank Zaw Min Htet and John P Gillies for help with protein purification, and Richard W Baker for help with cryo-EM data collection. We also thank the UC San Diego Cryo-EM facility, and the UC San Diego Physics Computing Facility for IT support. This paper was typeset with the bioRxiv word template by @Chrelli: www.github.com/chrelli/bioRxiv-word-template

## Materials and Methods

### Protein purification

Human dynein and Lis1 were purified as previously described (Htet et al., 2020).

### EM sample preparation

Grids were prepared as previously described (Htet et al., 2020). Briefly, UltraAuFoil R1.2/1.3 300 mesh grids (Electron Microscopy Sciences) were glow discharged with 20 mA negative current for 30 s. A 4 μL sample of 3.5 uM dynein monomer, 3.5 uM HaloTag-Lis1 and 2.5 mM ATP-VO_4_ was applied to the grid and vitrified using a Vitrobot Mark IV robot (FEI) set at 100% humidity and 4°C.

### EM data collection

Data collection was performed as previously described (Htet et al., 2020). Three datasets were collected and initially processed separately. Briefly, each dataset was processed in cryoSPARC using the patch motion correction and patch CTF extraction jobs to align micrographs and perform CTF estimation, respectively. Micrographs that had a CTF of >5 Å were discarded. Dose weighted images were used for particle picking using the crY-OLO training model generated in (Gillies et al., 2022; Wagner et al., 2019). Particles were extracted with a 1.16 Å/pixel. Several rounds of 2D classification were carried out in cryoSPARC to remove bad particles. Particles belonging to good 2D class averages in datasets 1 and 2 were combined and ab initio reconstruction was carried out. Particles belonging to dynein were combined and heterogenous refinement was carried out to separate intact dynein from partially unfolded dynein. Another round of heterogenous refinement was carried out that included the good particles from Dataset 3. Particles were separated into 1 Lis1 and 2 Lis1 classes, and each resulting map was used in nonuniform refinement (Punjani et al., 2020). The final resolution of human dynein-Lis1 and human dynein-(Lis1)2 was 4.0 Å and 4.1 Å, respectively.

We note that the overall resolution of our structures was limited due to preferred orientation. These datasets were collected on open hole grids before we began using streptavidin affinity grids, which helped reduce this problem in our most recent structure of yeast dynein-Pac1.

### Model building

The structure of Phi dynein (PDB 5NUG) and the AlphaFold model of human Lis1 (model P43034) were used as initial models for the human dynein-Lis1 structure and fit into the map using UCSF ChimeraX (Pettersen et al., 2021). Refinement of the model was carried out using a combination of Phenix Real Space Refine (Afonine et al., 2018) and Rosetta Relax (ver 3.13). Parts of the model were manually rebuilt using COOT (Emsley et al., 2010). Following completion of the human dynein-Lis1 model, it was used as a starting model for dynein-(Lis1)2 where Lis1_ring_ was duplicated and fit into the position at site_stalk_ using UCSF ChimeraX. Refinement proceeded using the same method as for dynein-Lis1.

## Video 1. Comparison of the human dynein-(LIS1)_2_ and yeast dynein-(Pac1)_2_ structures

The video compares the human (dynein-(LIS1)_2_) and yeast (dynein-(Pac1)_2_; PDB 7MGM) structures, highlighting some of the major interactions, and the differences in the positions adopted by LIS1/Pac1 at site_ring_ and site_stalk_ in the two systems.

## Video 2. Disease mutations in dynein and LIS1

This video shows the location of amino acids in LIS1 mutated in type-1 lissencephaly, and residues in dynein that are mutated in several neurodevelopmental or nondegenerative disorders (Charcot-Marie-Tooth, Spinal Muscular Atrophy, Autism Spectrum Disorders, and Malformations of cortical development/ Intellectual disability). We only show residues where we determined that the reported mutation(s) do not have an obvious destabilizing effect based on an inspection of the structure.

**Figure 1- figure supplemental 1.**
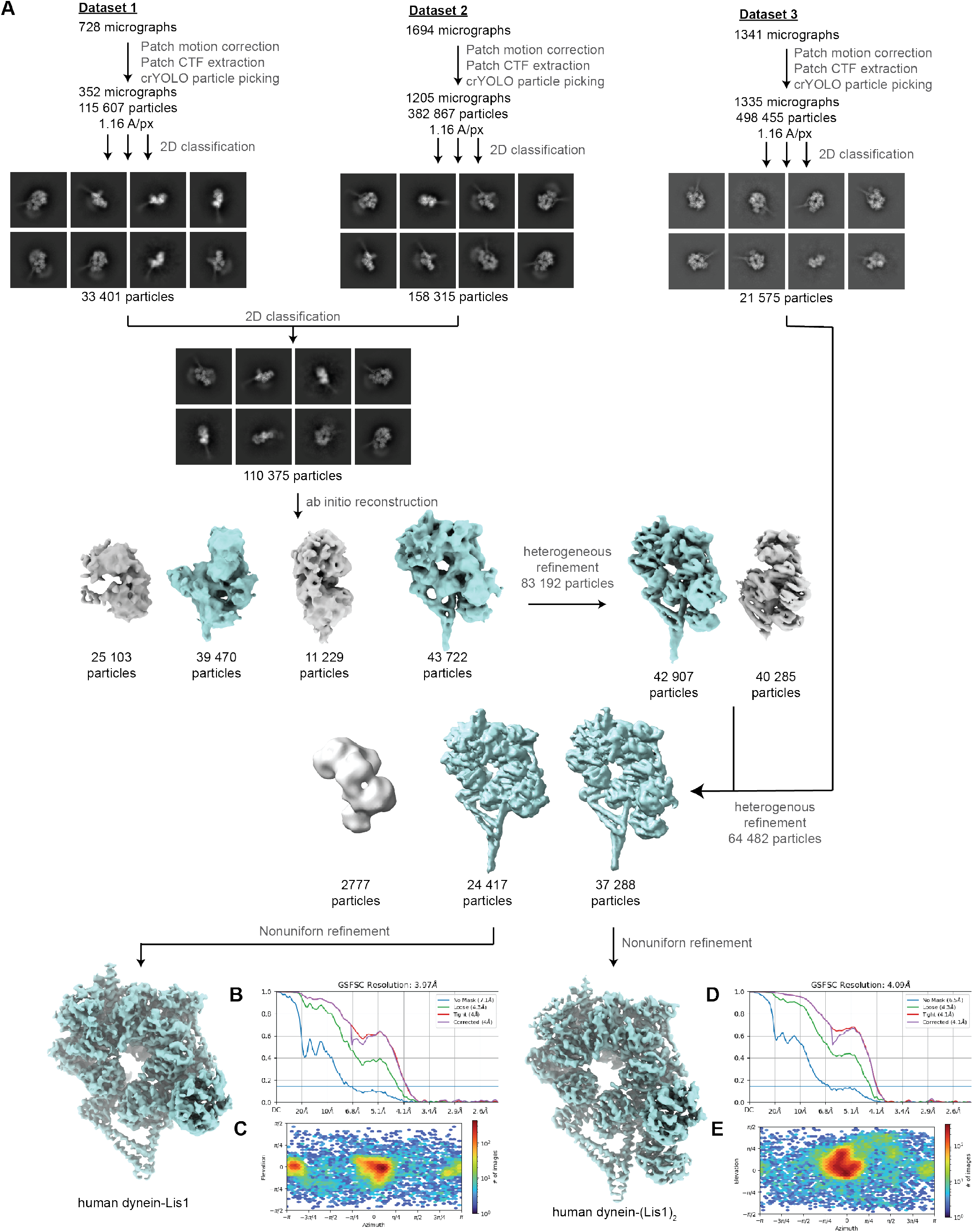
Cryo-EM data processing workflow. **(A)** CryoEM data processing pipeline for human dynein bound to LIS1. Particle extraction was carried out in Relion 3.0 and all other jobs were implanted in cryoSPARC. **(B, C)** FSC and angular distribution plots for human dynein-LIS1 (one β-propeller). **(D, E)** FSC and angular distribution plots for human dynein-(LIS1)_2_ (two β-propellers).

**Figure 1 – figure supplemental 2.**
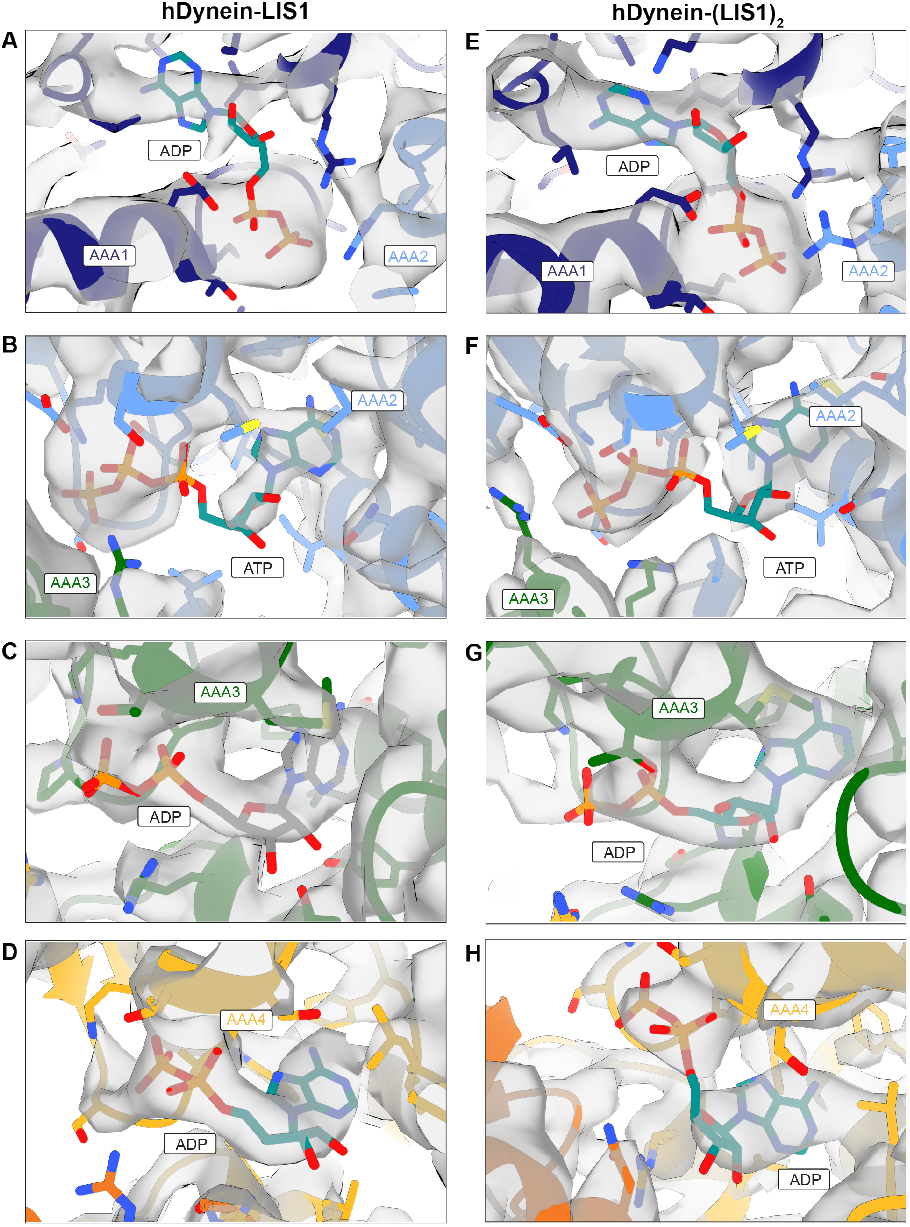
Nucleotides bound to dynein. **(A-D)** Nucleotides bound to dynein in the dynein-LIS1 (one β-propeller) structure at AAA1 (A), AAA2 (B), AAA3 (C) and AAA4 (D). **(E-H)** Nucleotides bound to dynein in the dynein-(LIS1)_2_ (twp β-propellers) structure at AAA1 (E), AAA2 (F), AAA3 (G) and AAA4 (H).

**Supplemental Table 1:**
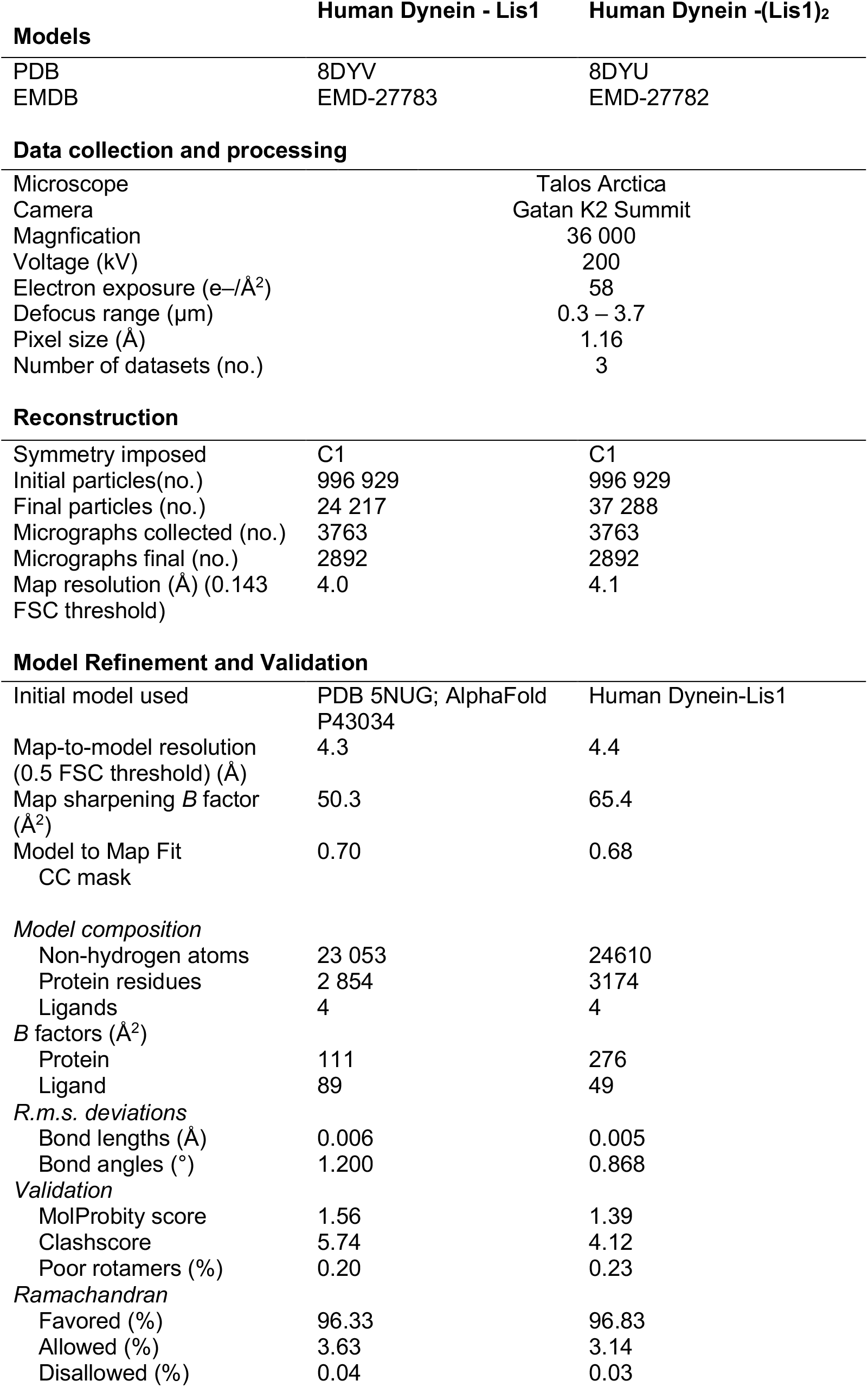
Cryo-EM data information and model validation. CryoEM data collection parameters, reconstruction information and model refinement statistics for the structures of human dynein bound to one and two LIS1 β-propellers.

